# Cross-Category Invariant Neurons in the Human Amygdala

**DOI:** 10.64898/2026.01.29.702502

**Authors:** Anna Bertucci, Stefania Picciallo, Olga Dal Monte, Marco Lanzilotto

## Abstract

Recognizing individuals or objects across different contexts is a hallmark of primate cognition. Beyond recognition, the brain organizes visual stimuli into meaningful categories, supporting efficient perception and adaptive behavior. Previous studies in the human temporal lobe have identified *concept cells* supporting invariant representations of specific stimuli. However, invariant coding alone cannot account for how categorical knowledge is structured across related visual domains. Here, we analyzed single-neuron recordings from the human amygdala during the presentation of familiar visual stimuli spanning multiple categories. Using a novel data-driven analytic framework that integrates supervised population decoding with single-neuron analyses, we identified a population of *cross-category invariant* neurons. These neurons preserved exemplar-invariant responses while generalizing across multiple categories linked by shared naturalistic or human/domestic contexts. Our findings demonstrate that the human amygdala supports both invariant and associative forms of categorical representations. By linking stable identity representations across contextually related categories, the amygdala provides a flexible neural substrate for high-level visual organization, supporting efficient perception and adaptive behavior in complex environments.

**Highlights:**

- Human amygdala neurons encode invariant category representations at the single-cell level
- Category representations are organized according to naturalistic and social contexts
- Cross-category invariant neurons bridge concept-cell and mixed-selectivity coding
- Category encoding relies on opponent patterns of enhancement and suppression across domain

## INTRODUCTION

Recognizing individuals or objects across different contexts is a hallmark of primate cognition. Beyond recognition, the brain groups visually distinct stimuli (such as an apple and a banana) into meaningful categories (e.g., fruit), thereby supporting efficient perception and adaptive behavior. This ability, highly constrained^1^ and evolutionarily conserved across species^2-4^, allows organisms to generalize across exemplars, prioritize behaviorally relevant information, and overcome major computational challenges.

Pioneering single-neuron recordings in the human medial temporal lobe (MTL) and inferotemporal cortex revealed neurons that respond invariantly to specific faces, individuals, animals or objects across variations in view, size, and context^5-9^. These discoveries established that single neurons—so-called *concept cells*—can encode stable, high-level representations independently of low-level visual features, supporting memory, recognition, and efficient identification of familiar stimuli^10,11^. Comparable evidence in macaques has demonstrated invariant coding for complex visual stimuli such as faces, objects and body postures^12-14^, and, remarkably, also for observed dynamic actions^15^, highlighting the computational challenge of maintaining stable identity signals across dynamic visual transformations^16^.

However, invariant coding alone cannot account for how the brain encodes the vast variability of the visual world. Flexible interaction with the environment requires organizing stimuli into *functionally meaningful* categories that prioritize ecologically and behaviorally relevant information while reducing computational demands. Although mixed selectivity—where single neurons respond across diverse contexts^17,18^—can enhance computational power and flexibility^19,20^, it does not by itself explain how meaningful associations emerge between related categories. It is therefore plausible that invariant coding and mixed selectivity operate as complementary mechanisms along a continuum, with individual neurons exhibiting intermediate properties that balance representational stability and contextual flexibility. How single neurons integrate invariant and context-dependent information remains a central question in systems neuroscience.

The amygdala, a limbic hub at the intersection of sensory, emotional, and associative processing, is ideally positioned to integrate visual identity with contextual relevance. Previous studies in both humans and non-human primates have reported selective amygdala responses to faces, objects, and social signals^13,21-23^, as well as a key flexible associative learning that links neutral stimuli to biologically significant outcomes^24^. Moreover, converging evidence suggests that neurons within the MTL contribute to the formation of long-term representations of stable associations, as defining features of human declarative memory^25^. Consistently, lesions or electrical stimulation of the amygdala can disrupt recognition or elicit vivid visual experiences, underscoring its role in forming adaptive, context-dependent categorical representations^26-28^.

Here, we directly address this gap by asking whether single neurons in the human amygdala encode purely invariant category representations or instead integrate stable identity signals with contextual associations across related categories. We analyzed a rare, large, and publicly available dataset of single-neuron recordings from the human amygdala^21^, acquired while participants viewed naturalistic images spanning familiar categories, including people, animals, and objects.

We identified a previously uncharacterized population of *cross-category invariant* neurons that exhibit stable, exemplar-invariant responses—resembling classical concept cells—while generalizing across multiple categories linked by shared contextual domains. These findings demonstrate that the human amygdala is not merely an emotional or salience detector, but a computational bridge between invariant and associative coding. By organizing visual information into ecologically and socially meaningful domains, the amygdala is ideally positioned to link perception to context-dependent action selection and physiological responses. More broadly, this work extends the classical concept-cell framework by uncovering an associative dimension of high-level representation within human limbic circuits, providing a mechanistic foundation for how perception, memory, and action-related processes are integrated into adaptive, context-sensitive behavior.

## RESULTS

We investigated whether the human amygdala forms contextual associations between meaningful categories by analyzing a large publicly available dataset^21^ of single-neuron recordings from 10 neurosurgical patients (**Table 1**). Participants viewed naturalistic images drawn from the Microsoft COCO database^29^, comprising 10 familiar categories (airplane, apple, bear, bird, car, chair, dog, elephant, person, zebra; **Figure 1A**), each with 50 unique exemplars per category. Neuronal activity was recorded while participants performed a one-back task (**Figure 1B**). On each trial, a single image was presented centrally following a brief fixation period (0.5-0.75-s). Participants were instructed to maintain fixation and passively view the images. In 10% of trials, randomly interleaved, the image was repeated, and participants were required to indicate whether the current image was identical to the preceding one (one-back repetition), in order to ensure task engagement. These one-back trials were excluded from all neural analyses to avoid potential motor or decision-related confounds. Each neuron was tested across the full stimulus set. Behavioral performance was reliable and well above chance, confirming task engagement (accuracy: 90% ± 14%; reaction time: 455 ± 28-ms; mean ± SD across subjects; **Figure 1C**).

**Table 1.**
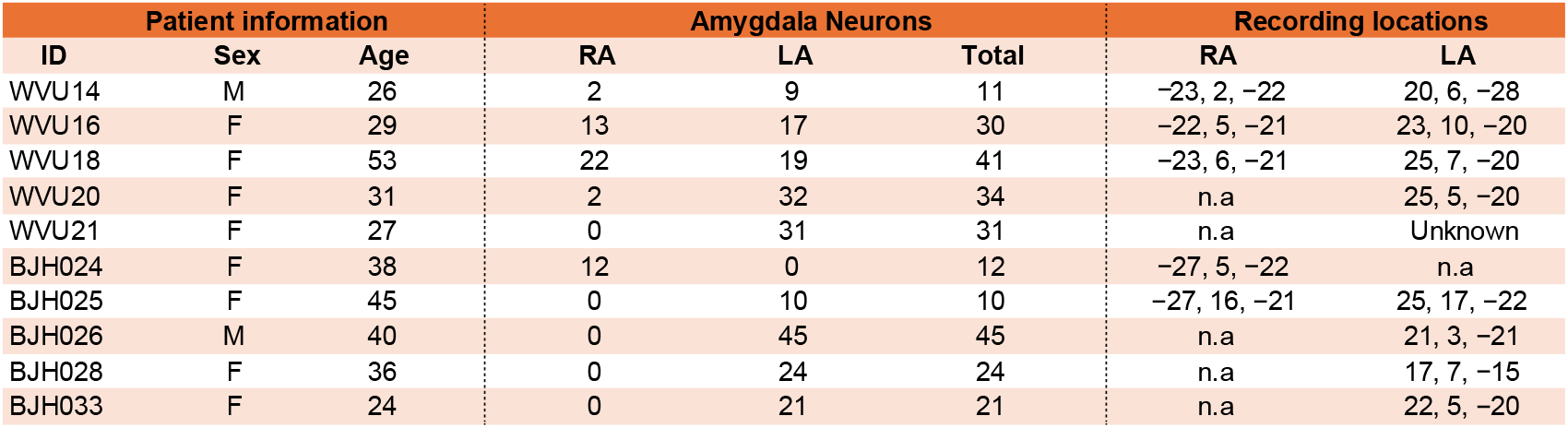
Patients and recording locations. Each row corresponds to a different patient. WVU indicates patients from West Virginia University and BJH indicates patients from Washington University in St. Louis. LA and RA indicate neurons recorded from the left and right amygdala, respectively. Radiological coordinates are reported in MNI space when available. Electrodes’ location was confirmed by the neurosurgeon. *n*.*a*., not available (no electrodes implanted). *Unknown*, imaging data is unavailable to determine precise electrode coordinates.

| Patient information |  |  | Amygdala Neurons |  |  | Recording locations |  |
| --- | --- | --- | --- | --- | --- | --- | --- |
| ID | Sex | Age | RA | LA | Total | RA | LA |
| WVU14 | M | 26 | 2 | 9 | 11 | -23, 2, -22 | 20, 6, -28 |
| WVU16 | F | 29 | 13 | 17 | 30 | -22, 5, -21 | 23, 10, -20 |
| WVU18 | F | 53 | 22 | 19 | 41 | -23, 6, -21 | 25, 7, -20 |
| WVU20 | F | 31 | 2 | 32 | 34 | n.a | 25, 5, -20 |
| WVU21 | F | 27 | 0 | 31 | 31 | n.a | Unknown |
| BJH024 | F | 38 | 12 | 0 | 12 | -27, 5, -22 | n.a |
| BJH025 | F | 45 | 0 | 10 | 10 | -27, 16, -21 | 25, 17, -22 |
| BJH026 | M | 40 | 0 | 45 | 45 | n.a | 21, 3, -21 |
| BJH028 | F | 36 | 0 | 24 | 24 | n.a | 17, 7, -15 |
| BJH033 | F | 24 | 0 | 21 | 21 | n.a | 22, 5, -20 |

**Figure 1.**
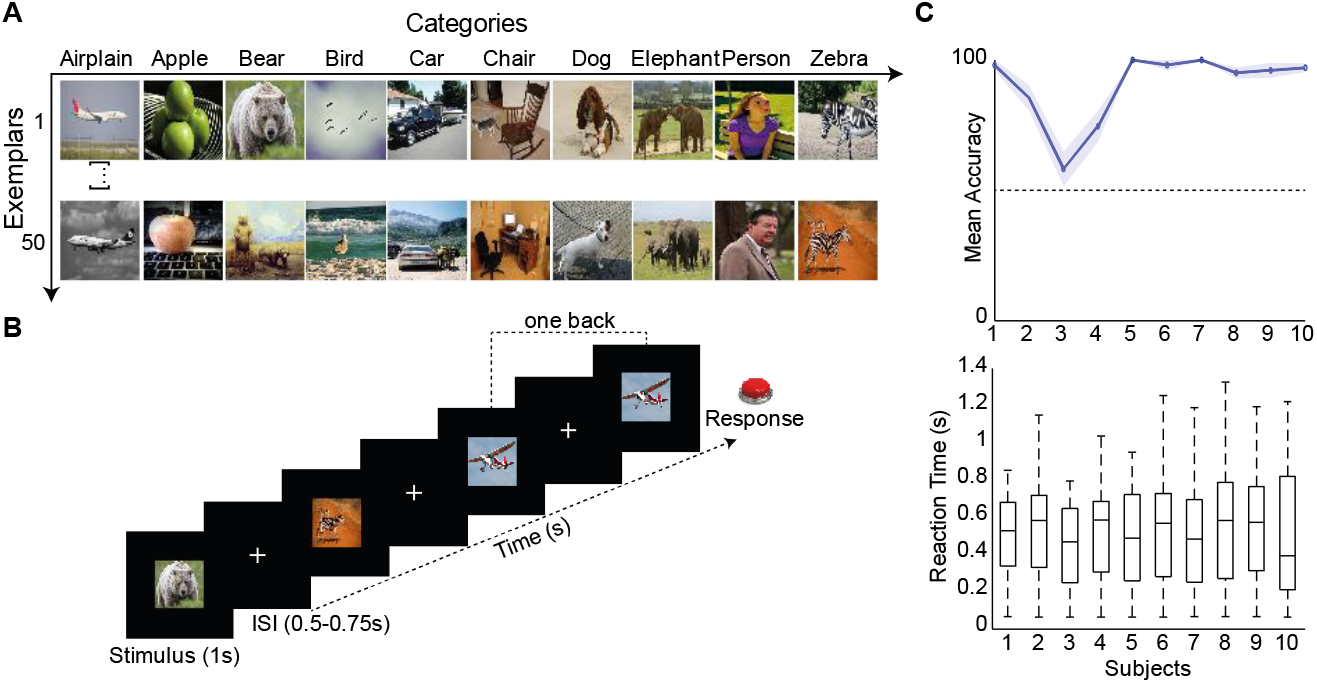
Experimental setup and behavioral performance. **(A)** Examples of stimuli from the Microsoft COCO dataset. Category labels are shown above each exemplar: *Airplane, Apple, Bear, Bird, Car, Chair, Dog, Elephant, Person*, and *Zebra*. **(B)** Behavioral Task. Participants performed a one-back fixation task in which they maintained fixation while images were presented at the center of the screen. In 10% of trials, participants were required to press a button when the currently presented image, randomly interleaved, was identical to the previous one. Each stimulus was displayed for 1 s, followed by an inter-stimulus interval (ISI) of 0.5-0.75s. Each image subtended approximately 10° of visual angle. **(C)** Behavioral performance in one-back trials. Mean accuracy was 90% ± 14% and mean reaction time (RT) was 455 ± 28-ms (mean ± SD across subjects). Top: line plot showing mean accuracy (range: 0–1) across participants for the 10% one-back trails; the gray shaded area indicates the standard deviation, and the dashed line marks chance-level performance (50%). Bottom: boxplots showing individual participants’ RTs during one-back trials, illustrating both within-subject variability and across-subject differences in RTs.

To characterize how category information is represented in the human amygdala, we applied a top-down analytic framework combining supervised population decoding, single-neuron analyses, and unsupervised clustering approaches. This strategy allowed us to identify data-driven neuronal classes encoding invariant category representations and contextual associations across related categories.

### Invariant population coding of familiar categories in the human amygdala

We analyzed 259 single neurons recorded from the amygdala of 10 neurosurgical patients across 12 recording sessions while they performed a one-back task (see STAR Methods and **Table 1**). To assess whether and how visual categories are represented, we applied a supervised population decoding approach using a Bayesian classifier^30^. Decoding performance (**Figure 2A**), evaluated by permutation testing^30^, was consistently above chance (*p* < 0.05), with peak accuracy remaining significant for a continuous 500-ms window. Category-related information emerged at ∼150-ms after stimulus onset and gradually declining thereafter. The square-shaped temporal profile of decoding accuracy indicates that category representations remained stable throughout stimulus presentation, consistent with the stationary nature of the exemplars. Although absolute decoding accuracy was modest, this outcome is expected given the low mean firing rates (1.73 ± 2.90 Hz; **Figure 2B**) typical of human amygdala neurons^21,31^, together with substantial trial-to-trial variability introduced by presenting only a single exemplar per category to preserve ecological validity (variability index = 0.78 ± 0.11; mean ± SD across subjects; **Figure 2C**). Both low firing rates and exemplar-driven variability are therefore expected to constrain classifier performance without implying weak or unreliable categorical encoding.

**Figure 2.**
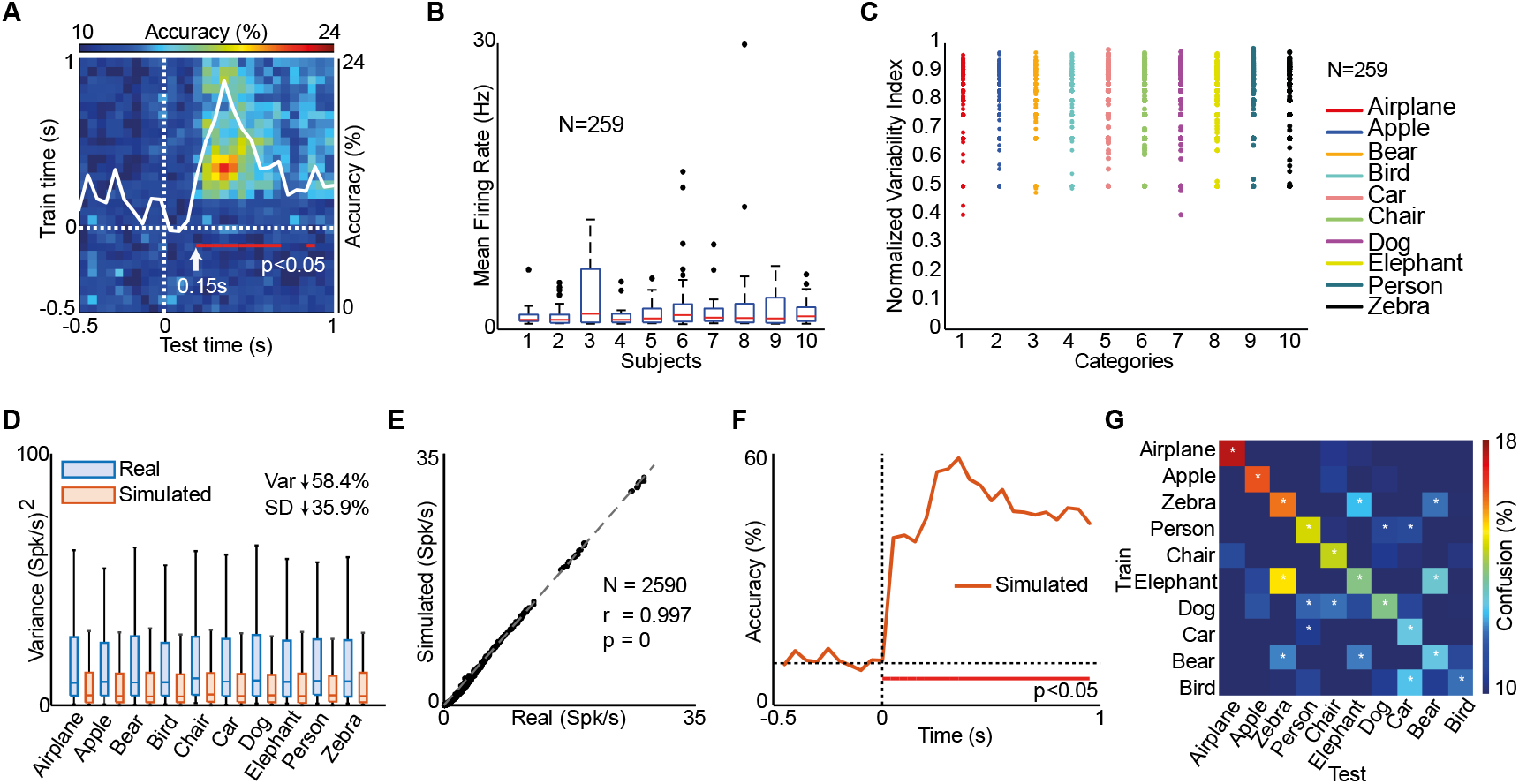
Population decoding of visual categories in the human amygdala and the impact of exemplars variability. **(A)** Time-resolved population decoding of stimulus category using a Bayesian classifier. Classification accuracy is shown as a function of training time (y-axis) and testing time (x-axis). The superimposed white trace represents the temporal evolution of decoding accuracy aligned to stimulus presentation. The red bar at the bottom indicates the time window during which decoding accuracy was significantly above chance (permutation test; *p* < 0.05; see STAR Methods). **(B)** Distribution of mean firing rates across neurons. Boxplots show firing rates (Hz) for each patient (N = 259 neurons total). Boxes indicate the interquartile range; red lines indicate the median, and black dots represent outliers. Overall firing rates were weak, consistent with typical human amygdala activity. **(C)** Trial-by-trial variability of neuronal responses. The scatter plot shows the normalized variability index (range 0–1) for each neuron across categories. Higher values indicate greater response variability across trials, reflecting exemplar-driven variability inherent to naturalistic stimuli. **(D)** Comparison of neuronal response variance between real data (light blue) and simulated data (light red). Simulated responses preserved the mean firing rate of each neuron for each category but reduced trial-by-trial variance by 58.4%, allowing direct assessment of the impact of variability on decoding performance. **(E)** Real versus simulated mean firing rates for all neuron–category combinations (2590 data points; r = 0.997, p < 0.001), demonstrating that the simulation preserved category-specific mean firing-rate tuning while selectively attenuating response variability. **(F)** Time course of category decoding accuracy for the simulated dataset (orange trace). The red bar indicates periods of decoding significantly above chance (*p*<0.05). Reducing trial-by-trial variability led to a marked increase in decoding accuracy, demonstrating that performance in real data is constrained by variability rather than weak categorical signals. **(G)** Confusion matrix summarizing category decoding performance during the 0.15–1-s post-stimulus window. Diagonal elements indicate correct classification, while off-diagonal values reflect systematic misclassifications. Errors were not randomly distributed but clustered within conceptually related categories (e.g., *Wild animals*: zebra–elephant–bear; *Domestic/Human-related*: person–chair– dog). White asterisks denote misclassification rates significantly higher than expected by chance (one-tailed permutation test; 1000 label shuffles; *p*<0.05), indicating structured associative relationships among categories.

To directly test this interpretation, we generated a synthetic dataset from original data in which trial-to-trial variability was reduced by ∼58.4% (**Figure 2D**), while preserving the mean firing rate of each neuron for each category (**Figure 2E**). Under these conditions, decoding accuracy increased markedly (**Figure 2F**). This manipulation demonstrates that classifier performance in the real data is primarily limited by exemplar-related variability rather than by the absence of categorical information in amygdala population responses. These results are consistent with previous single-neuron decoding studies^15^ under naturalistic conditions, which report comparable accuracy levels despite robust underlying representations.

Having established that amygdala population carries reliable information about category identity, we next asked whether this information reflects simple discrimination between categories or instead captures associative structure among them. To address this question, we trained a Bayesian discriminant classifier on all but one randomly left-out trial per category and evaluated performance on the held-out trials, summarizing results in a confusion matrix (**Figure 2G**). While some categories were reliably identified with minimal confusion (e.g., airplane, apple), misclassifications were not randomly distributed but instead clustered systematically within specific semantic groups, such as *Wild animal* (e.g., zebra, elephant, bear) or *Domestic/Human-related* categories (e.g., person, car, dog). A one-tail permutation test (1000 label shuffles; STAR Methods) confirmed that these structured confusion patterns were highly unlikely to arise by chance (*p* < 0.05).

Together, these findings demonstrate that the human amygdala supports not only invariant category identity but also encodes associative relationships among contextually related categories, reflecting naturalistic and human/domestic organizational principles.

### Cross-category invariant neurons: correlates of contextual generalization in the human amygdala

So far, we have shown that population activity in the human amygdala reliably encodes category identity and exhibits structured confusions among contextually related categories. We next asked whether this population-level organization arises from a small subset of highly stable and selective neurons or instead reflects broadly distributed mixed selectivity across the neuronal population.

To address this question, we applied an unsupervised category-preference ranking analysis^32^ to each recorded neuron (STAR Methods; **Figure 3A**). For each neuron, the preferred category was first identified during a reference epoch (250-750-ms post-stimulus onset). The rank of this preferred category was then tracked across time using sliding windows (200-ms width, 20-ms step) from baseline to stimulus offset. Time bins in which neural activity did not differ significantly from baseline were masked (white bins; sliding-window ANOVA, *p* > 0.05). Category preference stability was visualized using a color code (red: stable category preference; blue: unstable category preference). This analysis revealed marked heterogeneity in category preference stability across neurons (**Figure 3B)**. Some neurons exhibited highly stable tuning throughout stimulus presentation, whereas others showed rapid fluctuating preferences. For instance, Neuron 1 displayed strong tuning for airplanes, resembling classical concept cells. In contrast, Neuron 2 and 3 exhibited robust invariant responses that generalized across multiple categories linked by shared contextual domains (e.g., *Wild* and *Human/Domestic*). Notably, Neuron 3 showed response enhancement to *Domestic* categories and response suppression for *Wild* animal categories, as reflected in its net-activity profile. At the population level, 67/259 neurons (26%) met the criterion for stable category preference (≥120-ms of consecutive stable bins) during stimulus presentation, consistent with square-shaped temporal profile observed in population decoding.

**Figure 3.**
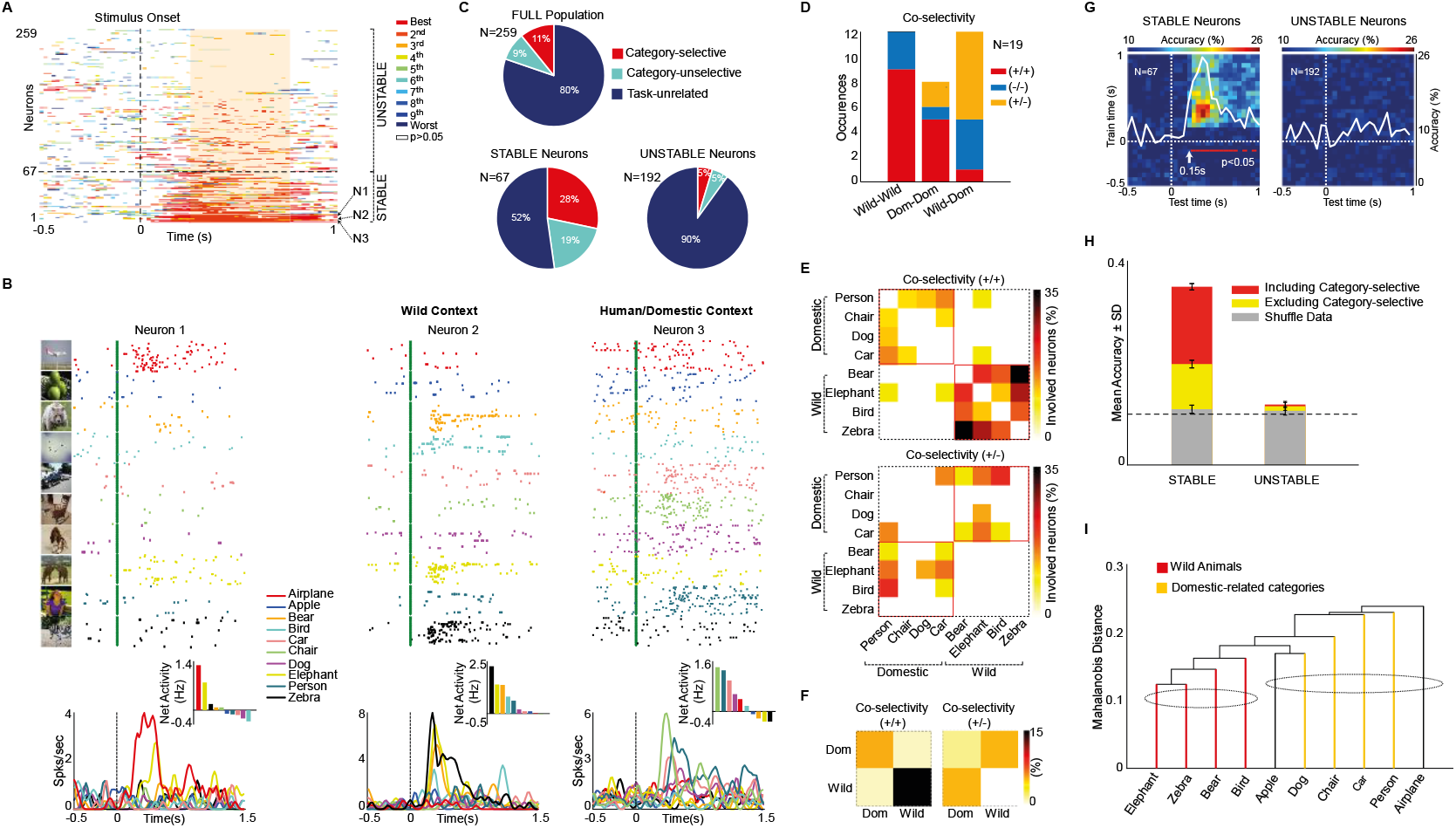
Temporal stability, cross-category selectivity, and contextual organization of amygdala neurons. **(A)** Category-preference ranking analysis for individual neurons. For each neuron, the preferred stimulus category (rank=1; *Best*) was defined during a reference epoch (250-750-ms after stimulus onset; shaded area) and its rank was tracked across time from the pre-stimulus baseline through stimulus offset using sliding windows (200-ms width, 20-ms step). Color indicates the instantaneous rank of the category identified as preferred in the reference window. Time bins in which neural activity did not differ significantly from baseline were masked in white (sliding-window ANOVA, 200-ms window, 20-ms step, *p*>0.05, uncorrected). Neurons were classified as *stable* or *unstable* based on the temporal persistence of category preference across consecutive time bins. N1, N2 and N3 indicate the relative position of example neurons (Neuron 1, 2 and 3) shown in B. **(B)** Example responses from three Category-selective amygdala neurons. Raster plots and peristimulus time histograms (PSTHs; 20-ms bins) are shown for all ten stimulus categories. Net firing-rate modulation (Hz) highlights enhancement (+) and suppression (-) effect across categories, computed over the 1 s period following stimulus onset. **(C)** Proportion of Category-selective, Category-unselective, and Task–unrelated neurons across the full population (top) and within stable and unstable subpopulations defined in (A). **(D)** Distribution of cross-category co-selectivity across neurons. Bars show the number of neurons exhibiting significant selectivity for more than one category, grouped as intra-context (Wild–Wild, Dom–Dom) or inter-context (Wild–Dom) pairs. Colors indicate the sign relationship between co-selective responses: enhancement–enhancement (+/+; red), suppression–suppression (−/−; blue), or opposite signs (+/− or −/+; yellow). **(E)** Population-level 8×8 matrix summarizing pairwise category relationships based on stable neurons. Each cell indicates the percentage of neurons involved in a given category pair that exhibit co-enhancement (+/+, top) or opposing modulation (+/–, bottom). Categories are ordered by contextual domain. Color intensity reflects the strength of co-selectivity. **(F)** Reduced 2×2 representation collapsing categories into two contextual domains (Wild vs. Domestic/Human-related). The matrix highlights predominant within-domain associations and systematic opposition across domains. Color scales as in (E). **(G)** Time-resolved population decoding of stimulus categories using stable (N=67) and unstable (N=192) neurons. Decoding conventions are as in Figure 2A. **(H)** Mean decoding accuracy (± SD) for stable and unstable neuronal populations. Bars show decoding performance in the 150-650ms post-stimulus interval using all neurons (red) and after exclusion of category-selective neurons (yellow). The gray bar indicates decoding accuracy obtained from shuffled labels. The dashed line denotes chance level (10%). **(I)** Hierarchical clustering of stimulus categories based on population responses of stable neurons performed in the 0.25-1 s post-stimulus interval, revealing two main branches corresponding to *Wild* (red) and *Domestic/Human*-related (yellow) contextual domains.

To further characterize these neurons, we performed a 3 (Epochs) x 10 (Categories) repeated-measure ANOVA followed by Fischer’s LSD post-hoc tests (*p* < 0.05; see STAR Methods). Based on this analysis, neurons were classified as Category-selective (28/259, 11%, e.g., Neuron 1-3), Category-unselective (23/259, 9%), or Task-unrelated (208/259, 80%) (**Figure 3C**). Importantly, the stable subpopulation contained a significantly higher proportion of task-related neurons (Category-selective plus Category-unselective; 32/67, 48%) compared with the overall population (51/259, 20%; χ^2^ = 45.03, *p* < 0.001). Conversely, only a small fraction of unstable neurons were task-related neurons (20/192, 10.4%), consistent with their rapidly fluctuating preferences.

Crucially, most Category-selective neurons within the stable set (17/19, 89%) did not behave as narrowly tuned “concept cells” selective for a single category. Instead, they exhibited *cross-category invariant responses* spanning multiple naturalistic categories. To quantify this co-selectivity structure, we grouped categories into two contextual domains based on ecological and social relevance: *Wild* animals (“Bear”, “Bird”, “Elephant”, “Zebra”) and *Domestic/Human-related* categories (“Car”, “Chair”, “Dog”, “Person”). “Airplane” and “Apple” were excluded to avoid ambiguous assignments. Co-selectivity was defined as a neuron showing significant post-hoc effects (Fisher’s LSD) for more than one category. For each co-selective pair, we determined whether it occurred within the same domain (Wild-Wild, Dom-Dom; *intra-context*) or across domains (Wild-Dom; *inter-context*), and whether the corresponding effects were of the same sign (enhancement/enhancement or suppression/suppression; +/+ or -/-) or opposite sign (+/-; **Figure 3D**). *Wild-Wild* pairs were entirely consistent in sign (12/12, 100%), *Dom-Dom* pairs were largely consistent (6/8, 75%), whereas *Wild-Dom* pairs were dominated by opposite-sign effects (7/12, 58%). A contingency table analysis confirmed that this distribution was significantly biased (χ^2^ = 14.37, df = 4, *p* = 0.0062), indicating that amygdala neurons preferentially link categories within the same contextual domain while opposing those across different domains. This reveals a dichotomic response pattern in which enhancement and suppression encode contextual structure rather than individual category identity.

To further examine this structure, we computed an 8×8 heatmap of pairwise enhancement across categories (excluding Airplane and Apple), ordered by Euclidean distance (**Figure 3E**, top). This analysis revealed strong clustering of within-domain associations (Wild or Domestic) alongside systematic opposition between domains (**Figure 3E**, bottom). Collapsing categories into two superordinate domains yielded a compact 2×2 representation (**Figure 3F**), confirming the predominance of within-context associations and robust anti-correlations between Wild and Domestic categories.

Notably, neurons with modest but consistent category preferences—identified by the preference-ranking analysis but classified as Category-unselective or Task-unrelated by ANOVA—also contributed to population decoding. This indicates that network-level categorical information is not limited to neurons exhibiting large firing-rate modulations. To test whether population-level information primarily arises from the stable subset, independent of ANOVA-based classification, we performed supervised decoding separately on stable and unstable neurons (**Figure 3G**). Decoding based on stable neurons closely matched the performance of the full population (**Figure 2A**), whereas unstable neurons did not contribute substantially. Strikingly, decoding remained significantly above chance even after excluding Category-selective neurons from the stable set (**Figure 3H**), demonstrating that stable but weakly tuned neurons also carry categorical information.

Finally, we examined whether stable neurons clustered in multidimensional response space according to contextual domains. Hierarchical clustering revealed two main branches corresponding to *Wild* and *Domestic/Human*-related categories (**Figure 3I**).

Together, these results demonstrate that high-level categorical information in the human amygdala is supported by a stable network of neurons whose coordinated patterns of enhancement and suppression encode contextual relationships across categories. Rather than reflecting isolated concept cells or diffuse mixed selectivity, amygdala representations emerge from structured cross-category invariance organized around ecologically and socially meaningful domains.

## DISCUSSION

Using a novel data-driven conceptual framework, this study demonstrates that single neurons in the human amygdala encode not only invariant category identity—classically associated with concept cells^10^—but also structured associative relationships across complex, naturalistic visual categories. By leveraging rare single-neuron recordings from neurosurgical patients^21^, we were able to probe high-level categorical representations in the human brain with a level of mechanistic precision that is inaccessible to non-invasive approaches and difficult to infer from animal models alone. Our results reveal that amygdala neurons support a form of categorical coding that balances stability and contextual flexibility—two defining hallmarks of primate cognition^33^ and language-based categorization^34^.

At the population level, category information was reliably decodable from amygdala activity, emerging rapidly after stimulus onset (∼150-ms) and remaining stable throughout stimulus presentation (**Figure 2A**). Importantly, decoding errors were not randomly distributed but exhibited a clear internal structure: misclassifications clustered systematically within ecologically and socially meaningful groups, such as *Wild animals* and *Domestic/Human-related* categories (**Figure 2G**). This pattern indicates that population responses captured relational structure among categories, rather than merely discriminating isolated category labels. Notably, this contextual organization emerged directly from the neural data, without imposing any semantic grouping a priori, underscoring its intrinsic nature.

This population-level organization was mirrored at the single-neuron level. By integrating temporal preference stability, classical selectivity analyses, population decoding, and hierarchical clustering (**Figure 3**), we identified a distinct subset of neurons—*cross-category invariant* neurons—that preserved exemplar-invariant responses while generalizing across multiple related categories. These neurons occupy an intermediate position between narrowly tuned concept cells^9-11^ and broadly mixed-selective neurons^18^, supporting the view that invariance and flexibility coexist along a continuum rather than constituting mutually exclusive coding strategies. Related principles have been reported in non-human primates, where parietal neurons encode observed actions invariantly across changes in the retinal input despite systematic rescaling of firing rate^15^, and in humans, where single neurons generalize action representations across perceptual and language-based descriptions^35^. Here, we extended these principles to human amygdala, showing that mixed selectivity does not replace invariant coding but embeds stable identity representations within structured neuronal spaces, enabling flexible generalization while preserving categorical stability.

A key insight of this work is that cross-category encoding in the amygdala follows structured patterns of enhancement and suppression across contextual domains (**Figure 3D, E, F**). Individual neurons showed enhanced responses to categories within the same domain while suppressing responses to categories belonging to opposing domains. This dichotomic, opponency-based organization was consistently observed across single-neuron analyses, pairwise enhancement matrices, and unsupervised clustering of the representational space, arguing against analytical bias or low-level visual similarity as alternative explanations. Instead, these convergent results indicate that amygdala neurons encode a genuine contextual organization of visual categories. This organizational principle aligns well with prior findings in non-human primates, where amygdala neurons encode stimulus value (positive vs. negative) through opponent firing patterns that flexibly reverse following changes in valence^36^. Our findings extend this principle beyond value coding, showing that similar opponency-based mechanisms operate at the level of visual category representations, linking ecologically and socially related categories while differentiating competing contextual domains.

This opponency-based associative coding aligns well with the established role of the amygdala as a hub for associative learning and behavioral control. The amygdala is critically involved in classical conditioning and social learning^24,37,38^, where experience shapes relationships between stimuli according to their behavioral relevance. It is tightly interconnected with hippocampal and para-hippocampal regions^39^, positioning it at the interface between rapid associative processing and long-term memory systems. While neurons in the human MTL encode durable associations among autobiographical entities such as people and places^25^, our findings demonstrate that single neurons in the human amygdala encode contextual relationships directly within visual category representations, potentially with an emotional valence, even in the absence of explicit memory demands or personal relevance. This organization is well suited to the amygdala’s anatomical and functional position. The amygdala receives convergent inputs from temporal and frontal cortices^40^, as well as fast subcortical visual signals via the superior colliculus and pulvinar^24,41,42^, granting early access to salient information. Through its outputs to motor and autonomic systems, these associations can be rapidly translated into adaptive behavioral and physiological reactions.

From a computational perspective, encoding categories through domain-specific enhancement and cross-domain suppression provides an efficient mechanism for mapping visual stimuli onto context-dependent action and physiological states, thereby supporting flexible behavioral control. By organizing representations in an opponency-based manner, the amygdala can bias competing behavioral states—such as approach versus avoidance, or freezing versus fight-or-flight^43,44^—using limited neural resources. This architecture enables rapid, context-sensitive selection of adaptative responses, including generalized body reactions^45,46^ or facial expressions^23,47^, while preserving stable categorical representations. Opponency-based coding thus offers a computationally economical solution for integrating perception, context, and action in dynamic environments.

In summary, this work reveals that *cross-category invariant* neurons in the human amygdala implement an associative and opponent form of categorical coding that extends beyond classical concept-cell representations. Rather than encoding categories in isolation, amygdala neurons organize invariant visual identities within a structured relational space, linking ecologically and socially related categories through coordinated enhancement and separating competing domains through suppression. This opponency-based architecture positions the amygdala as a computational bridge between stable recognition and flexible contextual interpretation, enabling rapid generalization and adaptive behavioral selection with minimal neural resources. By uncovering this mechanism at the level of single human neurons, our findings provide a cellular foundation for how perception, memory, and action are integrated into coherent, context-sensitive representations that support flexible behavior in complex environments.

## RESOURCE AVAILABILITY

### Lead contact

Requests for further information and resources should be directed to the lead contact, Marco Lanzilotto.

### Materials availability

Materials generated in this study may be available from the Lead Contact with a completed material transfer agreement.

### Data and code availability

- This study analyzes previously published and publicly available data deposited on the Open Science Framework (OFS; see key Resources Table), including stimulus images, behavioral data, sorted single-neuron spike times; and raw electrophysiological time-series data.
- This study does not report new original code.
- Any additional information required to re-analyze the data reported in this paper is available from the Lead Contact upon reasonable request.

## ACKNOWLEDGMENTS

We thank the authors for making this valuable resource available to the research community. This study made use of *A human single-neuron dataset for object recognition* published by Cao and collaborators^21^, which is distributed under a CC BY-NC-ND 4.0 license. This work was partially supported by the Brain & Behavior Research Foundation (BBRF 2021 Young Investigator Grant; Grant ID:30604), the Italian Ministry of University and Research-MUR (PRIN 2022; Grant ID: 2022Z8CXEM), the BIAL Foundation (Grant ID: 403/24), and the University of Parma (FIL_2024_PROGETTI_B_LANZILOTTO) to Marco Lanzilotto.

## AUTHOR CONTRIBUTIONS

Conceptualization: A.B., O.D.M. and M.L.; methodology: A.B., S.P., O.D.M, M.L.; writing–original draft: A.B. and M.L.; writing – review & editing: A.B., S.P., O.D.M, M.L.; funding acquisition: M.L.; supervision: O.D.M. and M.L.

## DECLARATION OF GENERATIVE AI AND AI-ASSISTED TECHNOLOGIES IN THE MANUSCRIPT PREPARATION

During the preparation of this work, the authors used ChatGPT solely for language editing and improvement of manuscript fluency. After using this tool/service, the authors reviewed and edited the content as needed and take full responsibility for the content of the published article.

## STAR★METHODS

### KEY RESOURCES TABLE

| REAGENT OR RESOURCES | SOURCE | IDENTIFIER |
| --- | --- | --- |
| **Software and algorithms** |  |  |
| MATLAB version R2023b | Mathworks Inc | https://www.mathworks.com/products/matlab.html |
| Neural Decoding Toolbox | Meyers, E. (2013) | https://readout.info/ |
| Adobe Illustrator | Adobe | https://www.adobe.com/it/products/illustrator.html |
| **Data** |  |  |
| Neurophysiological and behavioral recordings | Cao et al. (2025) | https://doi.org/10.17605/OSF.IO/VH5KQ |

### EXPERIMENTAL MODEL AND STUDY PARTICIPANTS DETAILS

We analyzed single-neuron recordings from neurosurgical patients with pharmacologically intractable epilepsy, obtained from a publicly available dataset^21^. Patients were recruited at West Virginia University (WVU) and Washington University in St. Louis (WUSTL) and implanted with depth electrodes targeting the amygdala for clinical monitoring.

For the present study, we selected a well-defined subset of 10 patients (5 from WVU and 5 from WUSTL; **Table 1**) for whom complete behavioral and electrophysiological data were available during performance of a one-back task using naturalistic images from the Microsoft COCO dataset^29^. This restriction ensured homogeneity of task demands, stimulus statistics, and recording conditions across subjects.

Analysis was limited to neurons with a mean firing rate > 0.15 Hz, a conservative threshold commonly used in human single-neuron studies to ensure reliable spike estimation while minimizing selection bias. This yielded a total of 259 well-isolated single units recorded from the amygdala. Each neuron was tested across the full stimulus set within a single recording session.

All participants provided written informed consent to participate in cognitive tasks during intracranial recordings and to allow sharing of de-identified data. Experimental procedures were approved by the Institutional Review Boards of West Virginia University (protocol #1709745061) and Washington University in St. Louis (protocol #202201019).

## METHODS DETAILS

### Stimuli and behavioral paradigm

Stimuli were static naturalistic images drawn from the Microsoft COCO dataset^29^. For this study, 10 familiar visual categories (airplane, apple, bear, bird, car, chair, dog, elephant, person, zebra) were selected with 50 unique exemplars per category (**Figure 1B**). Each image subtended approximately 10° of visual angle and was presented centrally.

Participants performed a one-back task designed to maintain visual attention while minimizing feature-specific strategies. On each trial, a single image was presented centrally for 1 s, followed by a uniformly jittered inter-stimulus interval (ISI) of 0.5–0.75-s (**Figure 1C**). Participants were instructed to maintain fixation and passively view the images. In 10% of trials, the image was immediately repeated, and participants were required to press a button when the current image was identical to the one presented on the preceding trial (one-back repetition), ensuring sustained task engagement. Each image was presented once, except when repeated as a one-back target. To avoid confounds related to motor preparation, decision-making, or repetition-related effects, all one-back repetition trials were excluded from neural analyses but were retained for behavioral performance assessment. This ensured equal sampling across images and categories: each neuron contributed responses to the full stimulus set, and the number of neuronal responses per image was matched across conditions.

### Data acquisition, electrophysiology, and spike sorting

All neural recordings, imaging co-registration, electrode localization, and spike sorting procedures were performed by the original authors and are described in detail elsewhere^21,31^. Here we summarize only the methodological aspects relevant to the present analyses.

Single-neuron activity was recorded from depth electrodes implanted in the amygdala of neurosurgical patients with pharmacologically intractable epilepsy. At each recording site, eight 40-μm microwires (Ad-Tech, WB09R-SP00X-014) were inserted into the clinical depth electrode. Bipolar wide-band signals (0.1– 9000 Hz) were acquired at a sampling rate of 32 kHz using Neuralynx systems (West Virginia University) or Blackrock systems (Washington University in St. Louis) and stored continuously for offline analysis.

Recording locations were estimated by co-registering post-implantation CT scans with pre-operative T1-weighted MRI scans and normalizing them to MNI space. Electrode positions were manually verified and confirmed by the neurosurgeon (see **Table 1**). Neurons recorded from seizure onset zones were excluded from all analyses.

Spike detection and sorting were performed offline by the original authors using a semi-automatic template-matching algorithm. Signals were band-pass filtered (300–3000 Hz), and spikes were detected using adaptive thresholding based on local noise estimates (typically >5× noise SD). Detected spikes were assigned to clusters based on waveform similarity, with continuous updating of cluster templates to accommodate electrode drift and short-term waveform variability.

Cluster quality was assessed using standard projection tests and isolation distance metrics^21^. The projection test quantified the separation between pairs of clusters recorded on the same channel by projecting spike waveforms onto the vector connecting cluster centroids. Isolation distance was computed as the Mahalanobis distance of the *n*th closest noise spike from the cluster center, where *n* corresponds to the number of spikes assigned to the cluster. Only well-isolated single units with a mean firing rate greater than 0.15 Hz and stable waveform and firing properties throughout the task were retained. Both broad-spiking and narrow-spiking neurons were included.

Recordings from different sessions of the same participant were obtained on separate days. To assess the possibility of recording the same neuron across sessions, the original authors performed control analyses comparing firing rate patterns and peristimulus time histograms across sessions. Similarity measures were not significantly higher within participants than across participants, indicating that neurons recorded across sessions were highly unlikely to correspond to the same single unit.

In the present study, we analyzed 259 well-isolated single neurons recorded from the amygdala during the one-back task with Microsoft COCO stimuli. No additional spike sorting, re-clustering, or cross-session unit merging was performed.

### QUANTIFICATION AND STATISTICAL ANALYSIS

All statistical analyses were performed using custom-made MATLAB scripts (MathWorks). Amygdala recordings were obtained across 12 recording sessions from 10 patients, yielding a total of 259 well-isolated single neurons.

#### Population decoding analysis

To assess population-level encoding of stimulus category, we employed a supervised population decoding framework (**Figures 2A, 2F**, and **3G**) based on a Poisson naïve Bayes classifier^15,30,32,48^, a generative model well suited for spike-count data. This approach quantifies how accurately neural population activity predicts the stimulus category presented on each trial, considering the activity of all recorded neurons jointly.

For each neuron, spike trains were binned into 150-ms windows sampled every 50-ms, yielding partially overlapping time bins. Within each bin, firing rates were computed by dividing spike counts by bin duration. At each time point, neural activity on a given trial was represented as a population vector of length *N* (number of neurons). For category decoding, population vectors were paired with their corresponding category labels (10 categories), with all trials contributing equally across conditions.

To obtain unbiased estimates of decoding performance, we used a balanced *k-fold cross-validation* procedure. Trials were partitioned into *k* folds (splits) such that each fold contained an equal number of trials from each category. On each iteration, the classifier was trained on *k – 1* folds and tested on the remaining folds. This procedure was repeated until each fold served as test data. To increase robustness, the entire cross-validation process was repeated 50 times with different random splits, and decoding accuracy was averaged across repetitions.

Statistical significance of decoding performance was assessed using a permutation test^15,30^. Category labels were randomly shuffled across trials, preserving the structure of population vectors while disrupting the relationship between neural activity and stimulus category. The decoding procedure was repeated 50 times on shuffled data to generate a null distribution. Decoding accuracy at each time point was considered significant (*p* < 0.05) when it exceeded all values in the null distribution.

To further characterize category-specific decoding behavior, we computed confusion matrices (**Figure 3G**) summarizing the proportion of correct and incorrect predictions for each true–predicted category pair. Classification accuracy within the matrix was quantified using the *zero–one loss metric* (fraction of correctly classified trials). Statistically significant misclassification patterns were identified using a one-tailed permutation test with 1000 label shuffles (*p* < 0.05).

#### Variability index

Trial-to-trial variability of neuronal firing was quantified using a soft normalized variability index (**Figure 2C**). For each neuron and for each of the ten stimulus categories, variability was computed as:

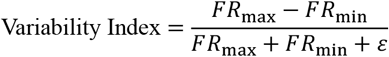

where *FR*_max_ and *FR*_min_ denote, respectively, the maximum and minimum firing rates observed across all trials of a given stimulus category for a given neuron. The constant ε = 1spike/s was added to the denominator to prevent (soft) division-by-zero and to reduce saturation effects for low-firing-rate neurons. This index ranges from 0 to 1, with higher values indicating greater trial-to-trial variability and lower values reflecting more consistent neuronal responses across repetitions. Variability indices were subsequently averaged across categories and neurons to obtain population-level estimates, reported as mean ± standard deviation (SD).

#### Category-preference ranking analysis

We applied an unsupervised category-preference ranking analysis to the full dataset of recorded neurons across subjects to quantify the temporal stability of category tuning^32^. For each neuron, we first identified the preferred stimulus category during a reference epoch defined as 250–750-ms after stimulus onset (0.5s window). This category was assigned rank = 1 and used as the reference category for that neuron. We then tracked the relative rank of this reference category across time using sliding windows (bin width: 200-ms; step: 20-ms) spanning from baseline (−0.5s before stimulus onset) to stimulus offset (1s after stimulus onset). Within each time bin, all ten stimulus categories were ranked according to the neuron’s firing rate, yielding a rank value from 1 (best; color code: red) to 10 (worst; color code blue) for the reference category.

Category preference dynamics were visualized using a color scale reflecting the rank of the reference category in each time bin. Time bins in which the reference category remained in the top-ranked category (rank = 1) were coded in red, indicating maximal preference stability. Progressive deviations from rank 1 were represented by intermediate colors, with bins in which the reference category dropped to lower ranks gradually shifting toward blue, indicating increasing instability of category preference.

To exclude time bins in which neural activity did not significantly differ from baseline, we applied a 2×1 repeated-measures *sliding-window* ANOVA (bin width: 200-ms; step: 20-ms), comparing firing rates between two epochs of each neuron: the pre-stimulus baseline (−0.5s) and the current sliding time bin. Time bins in which activity did not significantly differ from baseline (*p* > 0.05, uncorrected) were blanked out (color code: white).

Neurons were classified as *stable* if they exhibited at least 120-ms of consecutive time bins (i.e., ≥ 6 consecutive bins) in which the reference category remained ranked first (red bins). Neurons failing to meet this criterion were classified as *unstable*. This stability threshold was chosen to capture sustained category preference rather than transient or noisy fluctuations. This approach allowed us to dissociate stable and unstable category representations in a fully data-driven manner, independently of absolute firing-rate magnitude or predefined semantic category groupings.

### Classification of single neurons and relationship with preference stability

To characterize single neurons functional properties and relate classical selectivity to temporal preference stability, we combined the ranking-based stability analysis with a factorial ANOVA-based classification (**Figure 3C**). For each neuron, mean firing rates were computed for each of the 10 stimulus categories in two post-stimulus epochs: an *early* epoch (0–250-ms after stimulus onset) and a *late* epoch (250–1000-ms after stimulus onset), using a 0.5s pre-stimulus interval as baseline. A 3 (Epoch: baseline, early, late) × 10 (Category) repeated-measures ANOVA was performed (within-subject factors: *Epoch* and *Category*), followed by Fisher’s LSD post-hoc tests (*p* < 0.05).

Neural responses were classified as *enhancement* or *suppression* when mean firing rates in either post-stimulus epoch were significantly higher or lower than baseline, respectively. Based on these effects, neurons were assigned to one of three functional classes: 1) *Category-selective* neurons, showing category-specific modulation (post-hoc Category × Epoch effect) in at least one post-stimulus epoch; 2) *Category-unselective* neurons, showing significant task-related modulation relative to baseline without category specificity; 3) *Task-unrelated* neurons, showing no significant modulation across epochs or categories. When appropriate, Category-selective and Category-unselective neurons were grouped as *task-related* neurons.

Crucially, this ANOVA-based classification was orthogonal to the ranking analysis and served a complementary role. Whereas the ranking analysis quantified the temporal stability of category preference in a data-driven manner, independently of response magnitude, the ANOVA captured the statistical specificity and direction (enhancement vs. suppression) of neuronal responses. This combined approach allowed us to dissociate neurons that were temporally stable but weakly selective from those that were strongly selective but temporally unstable, and to identify a subset of neurons that jointly exhibited stable category preference and task-related modulation. These neurons constitute the core population of *cross-category invariant* neurons described in the Results, supporting the conclusion that temporal stability—rather than classical selectivity alone—is a key dimension of categorical coding in the human amygdala.

### Hierarchical cluster analysis

To further characterize the representational structure encoded by the stable neuronal population, we performed a hierarchical clustering analysis (**Figure 3I**) based on multivariate relationships among stimulus categories. For each of the ten categories, neuronal firing rates were computed in 20-ms time bins spanning the 250–1000-ms post-stimulus interval. Within each time bin, firing rates were averaged across trials (n=50) for each neuron, yielding population response vectors of size N=67 neurons. At each time bin, category-specific population responses were compared using a multivariate analysis of variance (MANOVA), and the resulting category centroids in population space were extracted. Pairwise distances between category centroids were then computed using Mahalanobis distance, with accounts for the covariance structure of the population responses. Categories were subsequently ordered from most similar to most dissimilar based on these distances. Hierarchical clustering was performed on the resulting distance matrix using a single-linkage criterion. The resulting dendrograms provide a data-driven visualization of the representational relationships among stimulus categories encoded by the stable neuronal population, without imposing any a priori semantic grouping.

